# Binding of a tetracationic *meso*-porphyrin to polyadenylic acid: a spectroscopic study

**DOI:** 10.1101/854380

**Authors:** Olga Ryazanova, Igor Voloshin, Victor Zozulya

## Abstract

Binding of a tetracationic porphyrin (TMPyP^4+^) to poly(rA) has been studied in neutral buffered solution of low ionic strength in a wide range of molar phosphate-to-dye ratios (*P/D*) using absorption spectroscopy, polarized fluorescence and fluorimetric titration. Two competitive binding modes were identified: partial intercalation of porphyrin chromophores between adenine bases prevailing at *P/D* > 20 and its outside binding to poly(rA) backbone dominating at *P/D* < 6. Both of them were accompanied by enhancement of the porphyrin emission. Absence of the emission quenching near stoichiometric *P/D* ratios allowed us to assume that external binding occurs without the self-stacking of the porphyrin chromophores.

## INTRODUCTION

Unrelenting interest to features of small molecules – nucleic acids (NA) interactions can be explained not only by the necessity to obtain their binding characteristics in the frame of fundamental science, but mainly by the possibility to apply the data obtained in biology, medicine, including synthesis of anticancer DNA-targeted drugs, as well as in nanotechnology [1–4]. In the last three decades, one of the most studied classes of such molecules are porphyrins, which belong to class of macrocyclic compounds with unique photophysical properties and the great potential in biomedical and molecular electronics applications [5–9]. Both spectroscopic properties and high biological activity of porphyrins are conditioned by the presence in its structure of planar macrocyclic chromophore [10], which effectively binds to DNA and RNA polymers, stabilizing and changing their local structure.

One of the most interesting porphyrin representatives, the water-soluble tetracyclic *meso*-porphyrin, TMPyP^4+^, is known as a fluorescent compound that binds effectively not only to double-stranded nucleic acids and to single-stranded, triple- and quadruple ones. Three main modes of TMPyP^4+^ binding to DNA have been established [11], which depend on the type of porphyrin substituents; on availability and type of a metal ion coordinated in the center of the tetrapyrrole macrocycle; on type, sequence and structure of the nucleic acid; on phosphate-to-dye ratio, *P/D*; ionic strength of the solution etc. There were indentified as intercalation, groove binding and outside binding without or with self-stacking. The last one can be accompanied by formation of the chiral porphyrin aggregates along the biopolymer chain [12,13].

It was shown that TMPyP^4+^ selectively accumulates in cancer cells, acts as efficient antitumor agent targeting G-quadruplex structures of telomeric DNA [14–18], and inhibiting telomerase activity with IC_50_ = 6.5 μM. It is widely used both in molecular biology as a probe for the structure and dynamics of nucleic acids [11,19,20], and in medicine as anti-viral and antimicrobal agent [21], photosensitizer for photodynamic therapy of cancer [22–27], and a carrier of antisense oligonucleotides for their delivery [28]. In complexes with redox-active transition metals (Mn^3+^, Fe^3+^, Cu^2+^, etc.) this porphyrin causes an oxidative DNA cleavage [28,30]. Molecular aggregates of TMPyP^4+^ with the chromophores stacking formed on NA exterior are characterized by the presence of strong electron interaction between adjacent porphyrin molecules. It determines their ability to transfer electron excitations and makes possible to use them in material science for the creation of new photonic materials as well as in molecular electronics for light-harvesting nanoelectronic devices [31–34].

The binding of TMPyP^4+^ porphyrin to nucleic acids has been most extensively studied on double-stranded [11,19,20,35,36] and quadruple DNA and RNA [37–39], whereas only limited number of publications were devoted to interaction of this porphyrin with single-stranded polynucleotides, including adenylate-containing homopolymers [12,40–43], despite the fact that single-stranded RNA and DNA segments are known to play important role in cell biology including replication, transcription and recombination processes [44,45]. For example, ss-poly(rA) was reported to be involved in regulation of the genes expression in eukaryotic cells [46]. It has been found in animal cells and many viruses in the form of sites up to 250 nucleotides in length, which are covalently bound to the 3’-end of polydisperse nuclear RNA and messenger RNA molecules [47]. Poly(rA) tails synthesized by poly(rA) polymerase (PAP) are known to play a significant role in the transcription process, as well as in stability and maturation of *m*-RNA [48]. Since PAP was found to be overexpressed in human cancer cells [49,50], ss-poly(A) segments can be suggested as a possible target for anticancer drugs – nucleic acid binders. Thereby compounds that selectively bind to the poly(rA) tail are supposed to be an inhibitor of the proteins synthesis in the tumor cells, as well as an effective target to control the function of cellular mRNA. Poly(dA) was proposed to be used in dynamic, DNA-based nanodevices as molecular switch, which at low pH forms a parallel-stranded double helix and at neutral pH exists as a ordered single helix [51].

Up to now the spectroscopic studies on interaction of TMPyP^4+^ with single-stranded adenine-containing homopolymers were carried out mainly on deoxyribonucleotide polymer poly(dA) [40, 12, 41], as well as oligomers (dA)_40_ [42] and (dA)_13_ [43] in solution of near-physiological ionic strength (0.1 and 0.2 M Na^+^). Whereas data on binding of this porphyrin or its metaloderivatives to ss-poly(rA) polyribonucleotide have been reported only in few works [41,12, 52]. At that the studies were performed mainly by absorption spectroscopy, CD and resonance light scattering (RLS) techniques. The changes in the porphyrin emission upon the binding to the polymer were followed only in [42,43]. Since fluorescent technique is known as a highly sensitive and informative tool, and binding of TMPyP^4+^ to anionic biopolymers was shown to be accompanied by significant changes in its emission properties [53–55,13], it seems appropriate to perform fluorescence study on binding of TMPyP^4+^ to poly(rA).

So, the aim of the current work is to study the binding of TMPyP^4+^ to ss-poly(rA) in aqueous solution of low ionic strength using polarized fluorescence and absorption techniques and to compare the results obtained with data obtained earlier for poly(dA) and poly(P).

## MATERIALS AND METHODS

### Chemicals

The tetra-*p*-tosylate salt of *meso*-tetrakis(4-N-methyl-pyridyl)porphine (TMPyP^4+^, Fig. 1) and potassium salt of poly(rA) from Sigma-Aldrich Chemical Co. were used without additional purification.

**Fig. 1.**
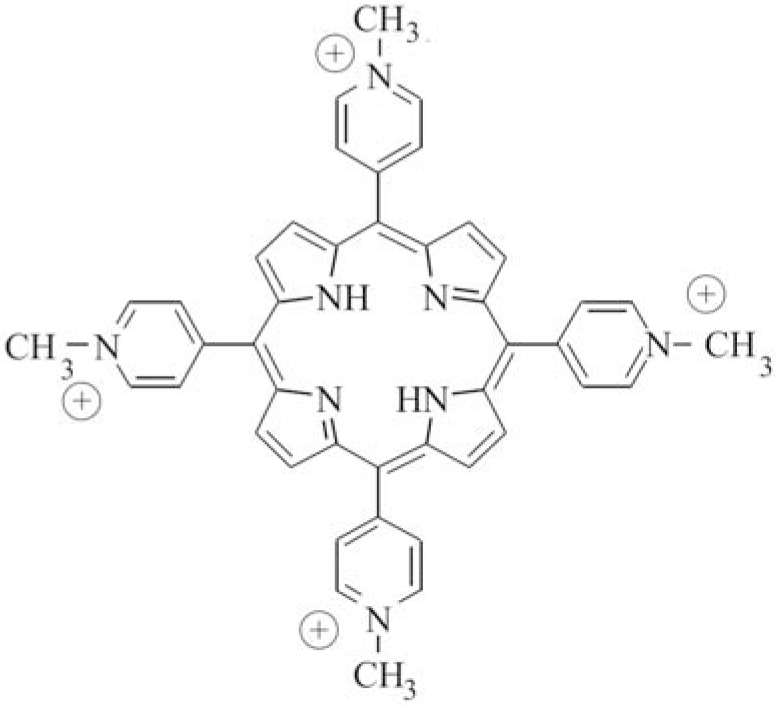
Molecular structure of tetracationic porphyrin TMPyP^4+^.

All experiments were performed in a 2 mM phosphate buffered solution (pH 6.9) prepared using the deionized water from Millipore-Q system. The porphyrin concentration was determined spectrophotometrically in water using the extinction coefficient of 226,000 M^−1^ cm^−1^ in the Soret band maximum [56]. It was 10 μM in all samples. The poly(rA) concentration was determined by UV spectrophotometry using the extinction coefficient ε_257_ = 10,100 M^−1^cm^−1^ (per mole of nucleotides, at 20 °C). [57].

### Apparatus and techniques

Electronic absorption spectra were recorded on a SPECORD UV/VIS spectrophotometer (Carl Zeiss, Jena).

Steady-state fluorescence measurements were carried out on a laboratory spectrofluorimeter based on the DFS-12 monochromator (LOMO, Russia, 350-800 nm range, dispersion 5 Å/mm) by the method of photon counting described earlier in [58]. The fluorescence was excited by stabilized radiation of a Philips halogen lamp in the region of the Q_y_(0,1) absorption band of TMPyP^4+^ porphyrin (λ_exc_ = 500 nm). The emission was observed at the 90° angle from the excitation beam. Ahrens prisms were used to polarize linearly the exciting beam as well as to analyze the fluorescence polarization. The spectrofluorimeter was equipped with a quartz depolarizing optical wedge to exclude the monochromator polarization-dependent response. Fluorescence spectra were corrected on the spectral sensitivity of the spectrofluorimeter.

The fluorescence intensity, *I*, and polarization degree, *p*, have been calculated using the formulas [59]:

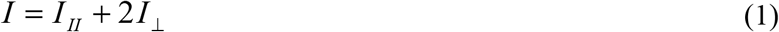

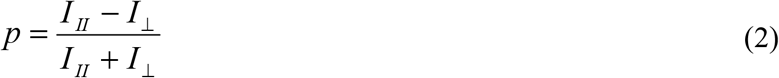

where *I*_*II*_ and *I*_⊥_ are intensities of the emitted light, which are polarized parallel and perpendicular to the polarization direction of exciting light, respectively.

Binding of TMPyP^4+^ to poly(rA) was studied in titration experiment where dependences of the dye fluorescence intensity and polarization degree at λ_obs_ = 670 nm on the molar phosphate-to-dye ratio, *P/D*, were measured under the constant dye concentration (10 μM). For this we add the dye sample with increasing amounts of the concentrated stock solution of poly(rA) containing the same dye content. *P/D* values were in the range from 0 to 2000. All experiments were carried out at ambient temperature (20 – 22 °C).

## RESULTS AND DISCUSSION

### Absorption and fluorescence spectra of free TMPyP^4+^ porphyrin

Visible absorption and fluorescence spectra of TMPyP^4+^ in a free state are presented in Figs. 2 and 3. It is seen, that the absorption spectrum (Fig. 2) consists of highly intense Soret band (or B-band) centered at 422 nm (ε = 226,000 M^−1^ cm^−1^ [56]) and four less intensive Q-bands in the region of 500-700 nm, corresponding to π→π^*^ transitions polarized within the plane of conjugated macrocycle. The porphyrin Soret band is known to results from superposition of two mutually perpendicularly polarized intense electron transitions [60] and corresponds to absorption of the photon energy from ground state (S_0_) to the second excited state (S_2_), whereas Q-bands (Q_y_(0,1) with maximum at 518 nm, Q_y_(0,0) at 551 nm, Q_x_(0,1) at 585 nm and Q_x_(0,0) at 641 nm [61]) relates to transitions from the ground state (S_0_) to the first excited one (S_1_). It is known that at studied porphyrin concentration (10 μM) the TMPyP^4+^ dimerization and self-aggregation can be neglected [62], so the spectrum of the free dye corresponds to its monomers.

**Fig. 2.**
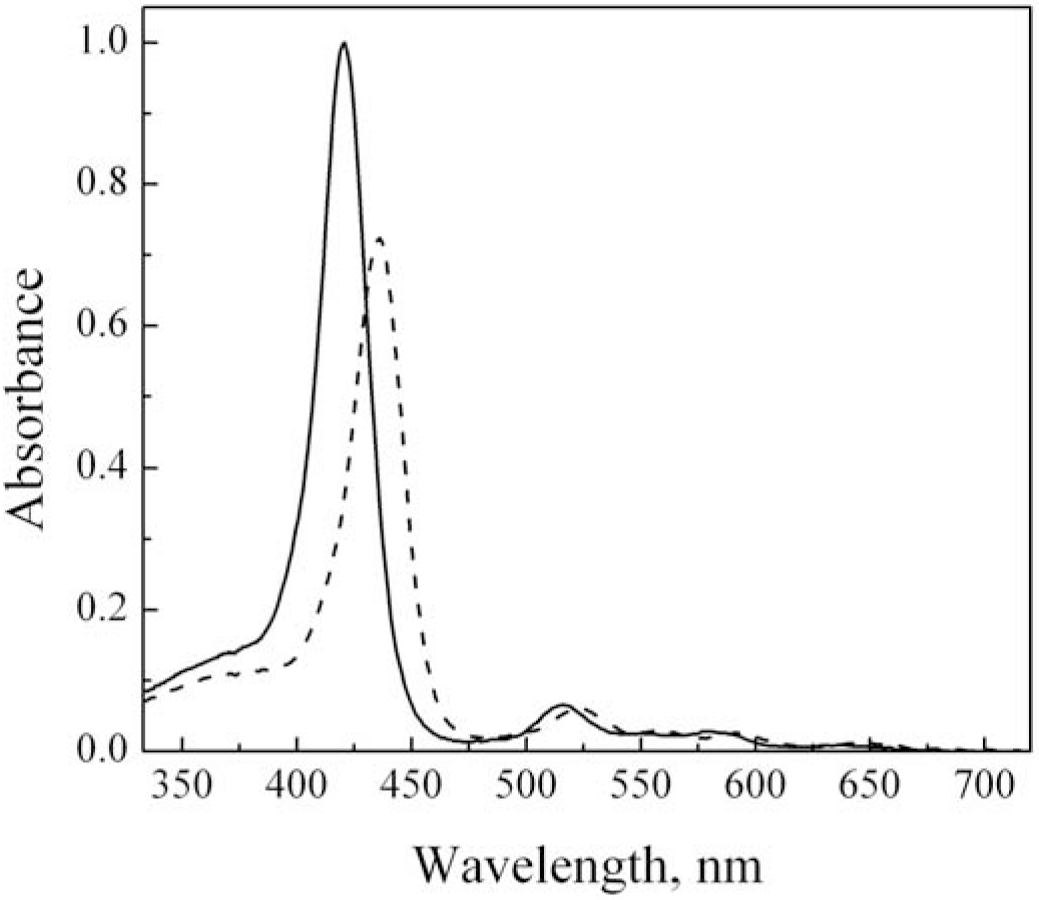
Absorption spectra of TMPyP^4+^ in a free state (solid line) and bound to poly(rA) at *P/D* = 1930 (dashed line) measured in 2 mM phosphate buffer, C_dye_ = 10 μM.

**Fig. 3.**
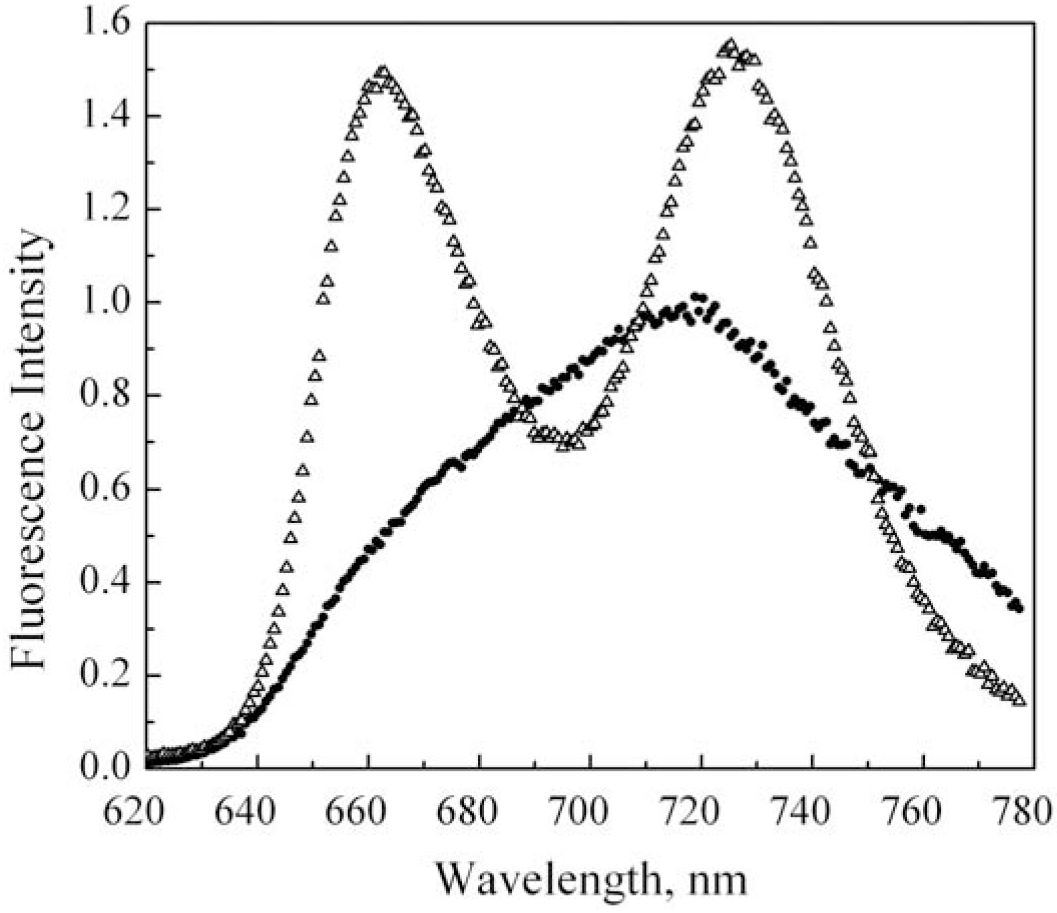
Fluorescence spectra of TMPyP^4+^ in a free state (●) and bound to poly(rA) at *P/D* = 1930 (Δ) in 2 mM phosphate buffer, C_dye_ = 10 μM, λ_exc_ = 500 nm is the wavelength of fluorescence excitation.

Emission spectrum of the TMPyP^4+^ (Fig. 3) is typical for cationic free-base porphyrins in water. It represents a broad asymmetric band with maximum at approximately 719 nm (for corrected spectrum, whereas for uncorrected spectrum λ_max_ = 678 nm) and characterizes by quite low value of fluorescence polarization degree (*p* = 0.02). Broadening of the spectra was explained by the result of two effects: intramolecular rotation of the porphyrin pyridinium groups and the mixing of the first excited singlet state of TMPyP4 with a close lying intramolecular charge transfer (CT) state involving partial transfer of charge from the core of the porphyrin to the electron-deficient pyridinium groups [62].

### Fluorimetric titration study of TMPyP^4+^ binding to poly(rA)

Fig. 4 show fluorimetric titration curves plotted as dependence of the relative fluorescence intensity, *I/I*_*0*_, and polarization degree, *p* (on inset), of TMPyP^4+^ measured at the fixed wavelength, 670 nm (near the maximum of uncorrected emission spectrum of free porphyrin), on molar *P/D* ratio. It is seen that titration curve is biphasic. In its initial range, corresponding to low *P/D* values (*P/D* < 6), addition of poly(rA) to the porphyrin solution results in rather steep enhancement of its emission intensity reaching *I/I*_*0*_ = 1.3 at *P/D* = 6, and fluorescence polarization degree from 0.02 (at *P/D* = 0) to 0.083 at *P/D* = 5.6 (Fig. 4, inset). With a further increase in the relative polymer content (at *P/D* > 15) the porphyrin emission continues to enhance but more gradually reaching *I/I*_*o*_ = 2.1 at P/D = 1200. The fluorescence polarization degree somewhat reduces and remains at a constant level of 0.075 (Table 1).

**Fig. 4.**
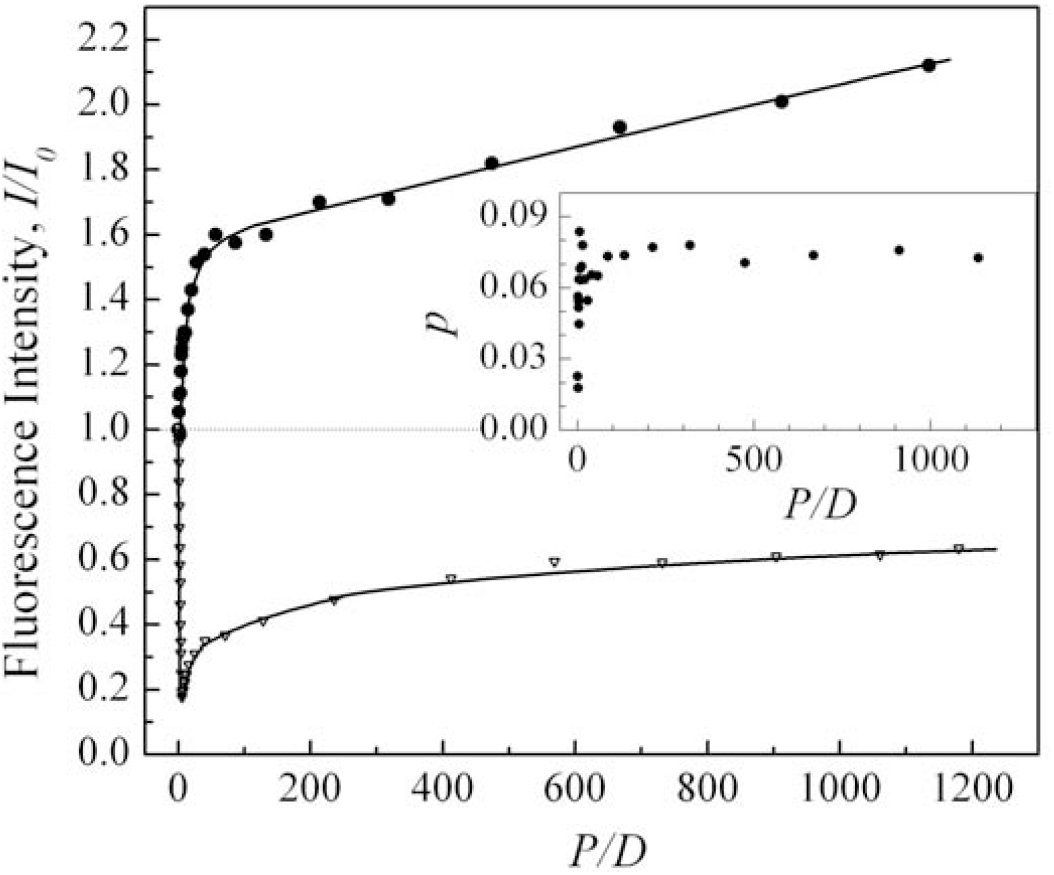
Fluorimetric titration curves plotted as dependence of the TMPyP^4+^ relative fluorescence intensity, *I/I*_*0*_, and fluorescence polarization degree (inset), *p*, on the molar phosphate-to-dye ratio, *P/D*, upon titration by poly(rA) (●). The data were obtained in 2 mM phosphate buffer pH 6.9 at constant porphyrin concentration [TMPyP^4+^] = 10 μM, λ_exc_ = 500 nm, λ_obs_= 670 nm. For comparison *I/I*_*0*_ *vs P/D* is presented for TMPyP^4+^ + poly(P) (∇) [54].

**Table 1.** Spectroscopic properties of TMPyP^4+^ porphyrin in a free state and bound to the biopolymers in 2 mM phosphate buffer, λ_exc_ = 500 nm

| Compound | Absorption <sup>a</sup> |  | Fluorescence |  |  |
| --- | --- | --- | --- | --- | --- |
| | $\lambda_{\text{max}}$, nm | $A/A_0$ | $\lambda_{\text{max}}$, nm | $I/I_0$ <sup>b</sup> | $p$ <sup>b</sup> |
| TMPyP<sup>4+</sup> | 422 | 1 | 719 | 1 | 0.020 |
| TMPyP<sup>4+</sup> + poly(rA)<br>($P/D = 5.6$) | – | – | – | 1.24 | 0.083 |
| TMPyP<sup>4+</sup> + poly(rA)<br>($P/D = 1930$) | 438 | 0.73 | 662.5, 724.7 | 2.18 | 0.075 |
| TMPyP<sup>4+</sup> + poly(P)<br>($P/D = 5.6$) | 410 | 0.49 | 667, 727 | 0.18 | 0.095 |
| TMPyP<sup>4+</sup> + poly(P)<br>($P/D = 1200$) | 420 | 0.77 | 715 | 0.63 | 0.034 |

<sup>a</sup> in the Soret absorption band <sup>b</sup> $\lambda_{\text{exc}} = 500$ nm, $\lambda_{\text{obs}} = 670$ nm for poly(rA) and 677 nm for poly(P)

### Absorption and fluorescence properties of TMPyP^4+^ bound to poly(rA) at high P/D ratios

Absorption and fluorescence spectra of TMPyP^4+^ bound to poly(rA) at high polymer content (*P/D* = 1930) are presented in Figs. 2, 3. It is seen that absorption spectrum exhibits 27% hypochromism and 16 nm bathochromic shift of the Soret band as compared with those for the free dye (Fig. 2). Substantial spectral transformation clearly indicates strong interaction between π-electronic systems of the porphyrin and nucleic bases. Fluorescence spectrum of the complex also undergoes significant changes (Fig. 3). Broad structureless emission band of the free dye (*P/D* = 0) upon binding to poly(rA) splits into the two well-resolved components with maxima at 663 and 725 nm that is accompanied by moderate enhancement of the porphyrin fluorescence intensity (*I/I*_*0*_ = 2.18) and polarization degree (*p* = 0.075) (Table 1).

## Discussion

From biphasic character of fluorimetric titration curve (Fig. 4) it is clearly seen that two binding modes are realized upon the TMPyP^4+^ interaction with poly(rA). The first one prevails at *P/D* < 6, whereas the second dominates at *P/D* > 20. Formation of the last type of complex at high *P/D* ratios is accompanied by substantial transformations of absorption and fluorescence spectra (Figs. 2,3). The observed 27% hypochromism and 16 nm bathochromic shift of the Soret absorption band (Table 1) clearly indicates strong interaction between π-electronic systems of the porphyrin and nucleic bases. A splitting of the wide featureless emission band of the free porphyrin into two components upon binding, as well as increase in its fluorescence intensity up to *I/I*_*0*_ = 2.18 evidences substantial changes in the chromophore microenvironment, namely the decrease of its polarity. Similar enhancement of the fluorescence intensity and splitting of the TMPyP^4+^ emission band was reported for the porphyrin solution in methanol, which has lower polarity than water [62]. Increase in the porphyrin fluorescence polarization degree upon binding (Fig. 4, inset) can be conditioned by deceleration of rotational motion of molecules, as well as increase in its excited state lifetime. From the foregoing it can be concluded that at high *P/D* ratios partial intercalation (or pseudointercalation) of porphyrin chromophore between neighboring adenine bases of poly(rA) is occurred. The magnitude of absorption changes observed in the present work at high *P/D* values (Table 1) are consistent with data reported for the same complex in solution of near-physiological ionic strength at *P/D* = 25 [40,41], where 34% hypochromism and 16 nm red shift of Soret band were observed. Authors proved pseudointercalation binding mode by characteristic changes in induced CD spectrum, namely by appearance of negative induced CD band at 457 nm.

By comparing the data for TMPyP^4+^ + poly(rA) at high *P/D* ratios with those obtained earlier for complex of TMPyP^4+^ with single-stranded poly(P) simulating phosphate backbone of nucleic acids [54], we can see that spectroscopic properties of porphyrin molecules partially intercalated between adenine bases and porphyrin monomers or dimers electrostatically bound to phosphate residues are absolutely different. So, for the last system 23 % hypochromism and 2 nm blue shift of absorption Soret band were reported, the emission band had broad featureless shape, and the fluorescence intensity is quenched up to 63 % from that for free dye (Table 1).

It is interesting, that binding of the TMPyP^4+^ to polydeoxyribonucleotide poly(dA) was accompanied by substantially less pronounced absorption changes then in the case of poly(rA). So, for the first system at *P/D* > 25 only 1 nm red shift and 14 % hypochromism of Soret absorption band were observed [41]. The difference can be explained by the significant distinction in the spatial structure of these two polynucleotides, including axial inter-base distance. So, ss-poly(rA) studied in current work at neutral pH represents right-handed 9-order A-type helix stabilized by the base stacking with pitch of 25.4 Å, so that the base planes are practically perpendicular to the helix axis [63–66]. Whereas poly(dA) relating to B-type helices is characterized by another conformation of sugar and has a more pronounced stacking of nucleic bases. The inter-base distance (axial rise per residue) for poly(rA) and poly(dA) was reported to be 2.8 Å and 3.3 Å correspondingly [63,65].

Another type of complexes formed between cationic porphyrins and anionic biopolymers and dominating at low *P/D* ratios is known as the electrostatic binding of the dye to the polymer exterior which can occur with or without self-stacking of the porphyrin chromophores. It have been revealed that near stoichiometric (on charge) *P/D* ratio this binding type can be accompanied by formation of the extended chiral porphyrin aggregates on the surface of the nucleic acids and other anionic biopolymers [12,54,56,67] that results in the absorption hypochromism, hypsochromic (for *H*-aggregates) or bathochromic (for *J*-aggregates) shift of the Soret absorption band, an appearance of strong resonant light scattering in the visible region and bisignate bands in induced CD spectrum (fingerprint of the excitonic interaction).

The nature of the binding and aggregation of porphyrin on the surface of nucleic acids and other polyanions was shown to depend significantly on the primary and secondary structure of the polymer. So, aggregation of TMPyP^4+^ with self-stacking have been revealed on the surface of ds-RNA [35], ordered single-stranded poly(rA) [12], poly(P) [54], pyrimidine homopolymers (dC)_40_ and (dT)_40_ (at *P/D* = 1.7) [42], however it was not found on ss-poly(dA) (investigation were performed at *P/D* = 2–25) [12], (dA)_40_, (dG)_40_ (at *P/D* = 1.7) [42]. As for native *ds*-DNA, the data available are controversial. So, aggregation of free-base TMPyP^4+^ have not been revealed on native DNA surface by Pasternack with co-aurhors [12] (investigation were performed in the range of *P/D* = 2–25), but it have been revealed in the work [13] where maximal aggregation was observed at *P/D* = 3.

Formation of TMPyP^4+^ self-assemblies on exterior of single stranded poly(dA) and poly(rA) have been studied earlier by Pasternack with co-authors [12]. Investigation were performed in 1 mM phosphate buffer containing 0.002–0.2 M NaCl, in *P/D* range from 0 to 40, using absorption, CD and RLS techniques. It has been revealed that poly(rA) promotes formation of the extended porphyrin aggregates bound to the polymer surface, and variations in drug load and ionic strength influence the tendency of porphyrins to self-assembly and affect the structure of the complexes formed. So, for TMPyP^4+^ + poly(rA) complexes in solution with 0.2 M NaCl formation of extended porphyrin aggregates on poly(rA) surface have been revealed upon *P/D* < 10. It was proved by appearance of enhanced resonance light scattering at 465 nm and characteristic large bisignate induced CD profile in Soret band region. In the range of *P/D* = 10 − 25 the large bound porphyrin aggregates were found, which at *P/D* > 25 disintegrate into the mixture of bound dye monomers and smaller aggregates. Variation of the solution ionic strength at constant *P/D* = 25 showed that aggregation of TMPyP^4+^ on poly(rA) exterior was occurred in the range of I = 0.002– 0.2 M NaCl, whereas for I > 0.2 it was not found. At the same time, no TMPyP^4+^ aggregation was found on poly(dA), that can be conditioned by substantial difference in the spatial structure of these polynucleotides. Authors have been concluded that proximity of anionic site and rigidity of the template backbone of poly(rA) as compared with poly(dA) are expected to encourage self-stacking of the cationic porphyrin molecules on its surface.

It is known that emission properties of TMPyP^4+^ are very sensitive to change in local environment. The binding of this porphyrin both to adenine nucleotide [67] and to adenosine 5’-monophosphate [55] was shown to results in the emission enhancement. At the same time formation of porphyrin dimers or aggregates with self-stacking of its chromophores results in the quenching of the dye fluorescence due to increase in the probability of non-radiative degradation of energy from an excited state [69]. The more perfect stacking between adjacent chromophores results in the stronger quenching of the porphyrin emission. So, formation of the TMPyP^4+^ aggregates on DNA exterior at near-stoichiometric binding condition was shown to result in the substantial quenching of the porphyrin emission [13], corresponding fluorimetric titration curve exhibited minimum at *P/D* = 3.2. In our work [54] it have been revealed that single-stranded inorganic polyphosphate, poly(P), representing linear flexible polyanion, can serve as a scaffold to assemble porphyrin molecules into weakly-fluoresces H-type aggregates. At near-stochiometric binding condition in the minimum of the fluorimetric titration curve complex of TMPyP^4+^ with poly(P) exhibited 53% hypochromism and 12 nm blue shift of the Sore absorption band, splitting of TMPyP^4+^ emission band into two components, the quenching of fluorescence up to 18% from the initial one, rise of fluorescence polarization degree and the appearance of strong resonance light scattering, which was maximal at *P/D* = 5.3 (Table 1). To compare behavior of TMPyP^4+^ upon outside binding to poly(rA) and poly(P) the fluorimetric titration curve have for the last polymer was added in Fig. 4. From the figure it is clearly seen that for TMPyP^4+^ + poly(P) the linear descending part of titration curve is observed in the *P/D* range from 0 to 6, whereas for TMPyP^4+^ + poly(rA) correspondent part of the curve has ascending character, emission intensity is increased up to *I/I_0_* = 1.3 at *P/D* = 6. Similar moderate enhancement of TMPyP^4+^ emission at low *P/D* ratios was observed upon its binding to oligodeoxyribonucleotides (dA)_13_ [43] and (dA)_40_ [42], where absence of porphyrin self-aggregation was proved by another techniques. In such a way, from absence of a fluorescence quenching at near-stochimetric (on charge) binding ratios and from the enhancement of the porphyrin emission observed in the range of *P/D* = 0 − 6 it can be concluded that external binding of TMPyP^4+^ to poly(rA) occurs without self-stacking of porphyrin chromophores.

## CONCLUSIONS

The present spectroscopic study shows that in neutral aqueous solution of low ionic strength (2mM Na^+^) water-soluble tetracationic *meso*-porphyrin TMPyP^4+^ effectively binds to single-stranded poly(rA). From biphasic shape of the fluorimetric titration curves two competitive binding modes are suggested: outside electrostatic binding of the porphyrin to phosphate backbone of the biopolymer prevailing at low *P/D* ratios (*P/D* < 6), and partial intercalation of porphyrin chromophore between adenine bases dominating at the polymer excess (*P/D* > 20). The last one provided by van der Waals’ forces and hydrophobic effect manifest itself by significant spectral transformations: 27% hypochromism and 16 nm red shift of Soret absorption band as compared with those for the free porphyrin, splitting of TMPyP^4+^ fluorescence band into two components, enhancement of its emission and increase in the fluorescence polarization degree. The absorption changes observed at high *P/D* for TMPyP^4+^ complexes with poly(rA) belonging to A-type helix are substantially more pronounced that those for poly(dA) belonging to B-type helix, where only 14 % hypochromism and 1 nm red shift of Soret band were reported [41], that can be conditioned by difference in the spatial structure of these polymers, in particular, by distinct axial inter-base distances.

The behavior of TMPyP^4+^ upon its outside binding to poly(rA) differs substantially from those reported for single-strand inorganic polyphosphate [54] and *ds*-DNA [13] where strong quenching of the dye fluorescence were observed at low *P/D* ratios as results of the formation of weakly fluorescing porphyrin aggregates with self-stacking. In the present work enhancement of the porphyrin emission was observed in all range of *P/D* studied. From absence of a fluorescence quenching upon fluorimetric titration at near-stochimetric (on charge) binding ratios it can be concluded that external binding of TMPyP^4+^ to poly(rA) occurs without self-stacking of the porphyrin chromophores.

Efficient binding of TMPyP^4+^ to poly(rA) may be the basis of its high biological activity and can be used for design of antitumor and antiviral pharmaceuticals. Also our findings provide new insight into the features of molecular interactions between macrocyclic dyes and nucleic acids.

## DECLARATION OF INTEREST STATEMENT

The authors declare that our manuscript complies with the all Ethical Rules applicable for this journal and that there are no conflicts of interests.

